# Shell Morphology, Radula and Genital Structures of New Invasive Giant African Land Snail Species, *Achatina fulica* Bowdich, 1822,*Achatina albopicta* E.A. Smith (1878) and *Achatina reticulata* Pfeiffer 1845 (Gastropoda:Achatinidae) in Southwest Nigeria

**DOI:** 10.1101/2019.12.16.877977

**Authors:** Alexander B. Odaibo, Suraj O. Olayinka

## Abstract

The aim of this study was to determine the differences in the shell, radula and genital structures of 3 new invasive species, *Achatina fulica* Bowdich, 1822,*Achatina albopicta* E.A. Smith (1878) and *Achatina reticulata* Pfeiffer, 1845 collected from southwestern Nigeria and to determine features that would be of importance in the identification of these invasive species in Nigeria. This is the first report of *Achatina albopicta* and *A. reticulata* in Nigeria, but *Achatina fulica* have since been reported in Nigeria and other African countries outside coastal East Africa. No study has described the external or internal morphology of any of the invasive species in Nigeria. Five to ten live specimens of each species, with complete shell characters, of each species were used for this study. Vernier caliper was used to obtain all shell measurements, with the shell held vertically and the aperture facing the observer. The genital structures were dissected out and fixed in 70% alcohol for 10-15 minutes and examined. The buccal mass was dissected out and digested in 7.5% sodium hydroxide for 24 hrs to free the radula from snail tissues and then examined under the compound microscope.

The shells of the 3 new species were dextral, conical with pointed spire and narrow apex. The whorls were separated by deep sutures. The parietal walls and the columella of the three species were white but columella of *A. reticulata* had a characteristic thick deposit of white porcelain-like material. There were dark brown markings on the whorls of the three species on dirty brown background for *A. fulica* and *A. reticulata* and dirty yellowish background for *A. albopicta*. The shell of *A. albopicta* was slightly glossy on the body whorl. The whorls of *A. albopicta* were much more convex than the whorls of *A. fulica and A. reticulata*. The columella of *A. albopicta* was truncate above the base of the peristome, moderately concave and slightly curved up at the base, while the columella of *A. fulica* was truncate sharply at the base of the peristome and straight and the columella of *A. reticulata* was slantly truncate at the base of the peristome and straight. The genitalia of the three species were very identical but differed slightly in the emergence of the basal vas deferens from the penis. The penes were slender and completely enclosed by the penial sheaths. The length of the penis varied from 10 to 12 mm. The vas deferens, free oviduct and the spermatheca duct were very long. The radula could be differentiated by the structure of central teeth and the first lateral tooth. The study showed that the shell morphology, radula and genital structures can be of importance in the identification of members of the family Achatinidae in Nigeria.

## Introduction

Species of Achatinidae are endemic in African countries south of the Sahara. They are characterized by large to medium sized, broadly ovate shells, with regular conical spires. With increase in mobility of humans and globalization of travel and trade, several achatinids, have been accidentally or purposefully transported to areas outside their native range, where they cause significant economic and ecological impacts [1]. *Achatina fulica* Bowdich, 1822 and its subspecies are native to the coastal East Africa, particularly Kenya and Tanzania but have now been introduced to many other African countries like Cote d’Ivoire, Togo, Ghana, Nigeria [2-6], Benin Republic were *A. fulica* has already overtaken the west African land giant snails in population density [7], and many countries in tropical and subtropical regions [1].

The Giant African land snails *Achatina fulica* Bowdich, 1822 is considered to be among the world’s 100 worst invasive species and also ranked among the worst snail pests of tropical and sub-tropical regions, causing significant damages to farms, commercial plantations and domestic gardens [8]. It is an intermediate host of the rat lungworm, *Angiostrongylus cantonensis*, which causes eosinophilic meningoencephalitis in humans and *Angiostrongylus costaricencis*, the etiological agent of abdominal angiostrogylosis [9, 10]. *A. fulica* can consume several species of native plants, agricultural and horticultural crops, modify habitats and outcompete native species [11]. In Nigeria the West African species of *Archachatina* and *Achatina* (subgenus *Achatina)* are usually favoured and they grow to reasonable sizes than any other known land snails. The genus *Achatina* had been represented by *Achatina achatina* (Linne) and its subspecies, but in recent times *Achatina fulica* (subgenus *Lissachatina)* has been reported in some parts of the country [5,6]. A recent study of the live specimens of *Achatina* spp collected from Itori, in Ogun state and the University of Ibadan, Ibadan, Nigeria in Oyo state, showed that the samples collected were represented by three separate species (*Achatina fulica, Achatina albopicta and A. reticulata*),following the original descriptions of Bequaert [12]. Shell morphology alone could not reveal clearly the differences between these species, hence a combination of shell, radula and genital structures known to be important in the classification molluscs [13] were investigated for their significance in the identification of the invasive snail species. This study was undertaken to reveal the differences in the shell, radula and genital structures of the three invasive species and provide notes on characters that can be important for their identification in areas where they have been recently introduced and to contribute to the available information on the terrestrial molluscan species richness in Nigeria that is still poorly investigated.

## Materials and Methods

### Snail samples

The samples of *Achatina fulica* used for this study were collected from Itori, headquarter of Ewekoro Local Government Area in Ogun State, Nigeria, located at 6°93’23” N and 3°22’47”E. The average monthly temperature ranges from 23°C to 32.2°C and it is a tropical rain forest area undergoing transition to guinea savannah due to farming and mining activities. The samples of *Achatina albopicta* and *A. reticulata* were collected from the residential areas of University of Ibadan, Ibadan, Oyo state, Nigeria, located at 7°23’28.19” and 3°54’59.99” E “Fig.1”. Five to ten live snails were used for the study. Snails were identified using Bequaert [12].

**Fig 1.**
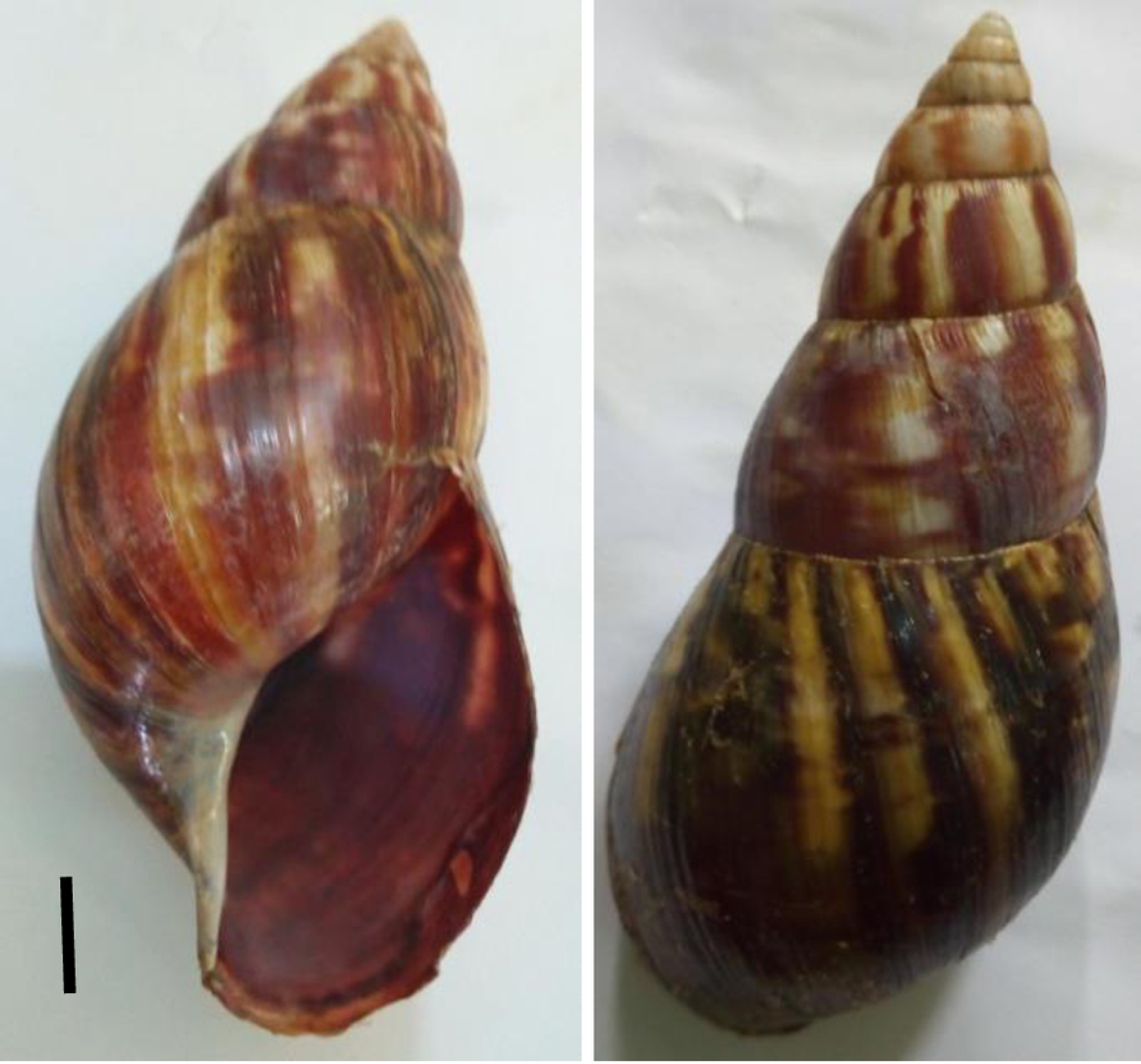
Map of Oyo and Ogun States Showing the Collection Sites for the Invasive *Achatina* Species in Nigeria.

### Snailery

Snail samples collected from the field were maintained in the snailery in the Department of Zoology, University of Ibadan, Ibadan, Nigeria, before examination. Snails were kept in rectangular glass containers (93 × 62 × 58 cm), with metal mesh covers, field half way with moist humus soil. The snails were maintained under the natural regime of 12hr light and 12hr darkness, fed *ad libitum* with *Carica papaya* leaves, *Tridax triangulare* leaves and unripe pawpaw fruits before examination.

### Shell morphometrics

Vernier caliper was used to obtain the conchological character measurements [14]. All measurements were taken with the shell held vertically with the aperture facing the observer, and only complete shells i.e. shells with no missing parts, were used for the study. The shell height (SH) was the longest vertical axis of the shell, measured from the tip of the spire to the basal edge of the outer lip. The shell width (SW) was the largest diameter, measured at right angle to the vertical axis, from the left margin of the body whorl to the outer edge of outer lip. The aperture height (AH) was the longest distance from the insertion of the outer lip on the parietal wall to the basal edge of the outer lip, while the aperture width (AW) was the greatest distance from the inner edge of the columella to the edge of the outer lip. The spire height (SpH) was measured from the tip of the apex to the suture separating the spiral whorls from the body whorl, and the body whorl height was the measured from the suture separating the spire from the body whorl to the base of the peristome (BwH). All measurements were in millimeters.

### Shell parameter indices

From the values obtained for each linear measurement, the following indices were determined, following the methodology of Medeiros et al.[15]: the shell height/shell width (SH/SW); body whorl height/shell width (BwH/SW); shell aperture height/shell aperture width (AH/AW); and spire height/ body whorl height (SpH/BwH).

### Number of whorls

The number of whorls corresponds to the number of complete turns of the shell beginning from the embryonal whorls at the tip of the spire.

### Dissection of genital structures

Each specimen was placed in a jar filled with boiling water for 10 – 15 minutes to loosen the columellar muscle and the soft body was extracted from the shell with hooked metal. The shell was preserved for shell morphometrics. The genital structure was dissected out of the soft body and fixed in 70% alcohol for 10 minutes and examined. The genital structure was then preserved in 10% formalin.

### Radula preparation

The buccal mass was dissected out of the head region of the soft body and macerated in 7.5% sodium hydroxide for 24 hours at room temperature. The freed radula was washed in water and any leftover tissues were removed under the dissecting microscope. The radula was mounted in glycerine and view under the compound microscope. After observation the radula was preserved in 70% alcohol. Photographs of radula were taken with UCMOS series microscope camera MU500 with Toupview 3.2 image software.

### Statistical analysis

To detect variations in shell morphology between species, the linear measurements and ratios were subjected to logarithmic transformation (*log*_10_) before using Analysis of Variance (ANOVA, p≤0.05).

## Results and Discussion

The three new species in this study, *Achatina fulica, A. albopicta*, and *A. reticulata* were collected from residential areas in Ibadan, an urban community and from farmlands in Itori, a semi urban community in southwest Nigeria. The invasive species are known to thrive in the presence of man, especially in urban sites and farms or disturbed areas [8], and they have superior competitive abilities over endemic species, whose distributions are often affected, due to the high fecundity and reproductive rates of the invasive species [11]. This invasion will significantly impact the species richness of land snails in Nigeria. It is therefore important that the extent of this invasion in Nigeria is monitored and properly managed before they assume pest status or dominance over the West Africa species of achatinidae that have served as cheap sources of protein for the rural populace and have not been incriminated in the transmission of any known parasitic infections of public health importance. The invasive *A. fulica* has achieved dominance in the achatinid communities in Ivory Coast, Ghana and central Benin Republic [1, 7]. While it is not precisely known how the invasive species got to Nigeria; it is most likely that they were introduced for economic reasons and subsequent uncontrolled uses may increase their ranges in future as they are moved around for sale, food or farming by humans. What should also be of concern, apart from the effect on ecology and biodiversity, is that the invasive species can act as intermediate hosts of Metastrongylidae nematodes, particularly now that *Angiostrongylus cantonensis* Cheg, 1935, which causes eosinophilic meningitis is spreading rapidly to many parts of the world [10].

The shell morphology of the three invasive species (*Achatina fulica, A. albopicta* and *A. reticulata*) in this study conformed to the original descriptions of the three species from East Africa [12]. The shells were dextral, conical with pointed spires and narrow apex. They had a minimum of 8 whorls and the whorls were separated by deep sutures. The parietal walls and columella of the three species were whitish. There were dark brown markings on the whorls of the three species, but on dirty brown background for *A. fulica* “Fig 2” and *A. reticulata* “Fig 3”, and dark brown or yellowish background for *A. albopicta* “Fig 4”, which was also slightly glossy on the body wall. The major differences in shell features are shown in “Table 1”.

**Table 1.** Showing differences in shell characters of the invasive Achatina species in southwestern Nigeria.

| Shell characters | *Achatina* species |  |  |
| --- | --- | --- | --- |
|  | *Achatina fulica* | *Achatina albopicta* | *Achatina reticulata* |
| Spire | The spire occupied 30% of the total height of the shell, with longitudinal brown stripe or patches. The whorls were slightly convex. | The spire occupied 32% of the total shell height, with light brown irregular patches or stripes. The whorls were strongly convex. | The spire occupied 29 % of the total shell height, with dirty white background and zigzag/ longitudinal brown patches or stripes. . The whorls were strongly convex. |
| Body whorl | Slightly glossy and terminates almost at the end of the columella. | Slightly glossy, and extends below the truncated end of the columella. | Very rough with longitudinal sculptures visible to the naked eyes. |
| Columella | White, nearly straight and sharply truncated at the base of the aperture. | White, concave, bent upwards and sharply truncated above the base of the aperture. | White porcelain-like deposit, straight and strongly truncated at a slant above the base of the aperture. |
| Parietal wall | Dark brown background with faint white | White with faint dark brown background | Thick white porcelain-like deposit |
| Inside of shell | White with bluish tinge on a dark brown background | White with bluish tinge | Very white with bluish tinge |
| Outer lip of peristome | Blackish brown and thin | Light brown and thin | Thick and white with broken brown patches |

**Fig 2.**
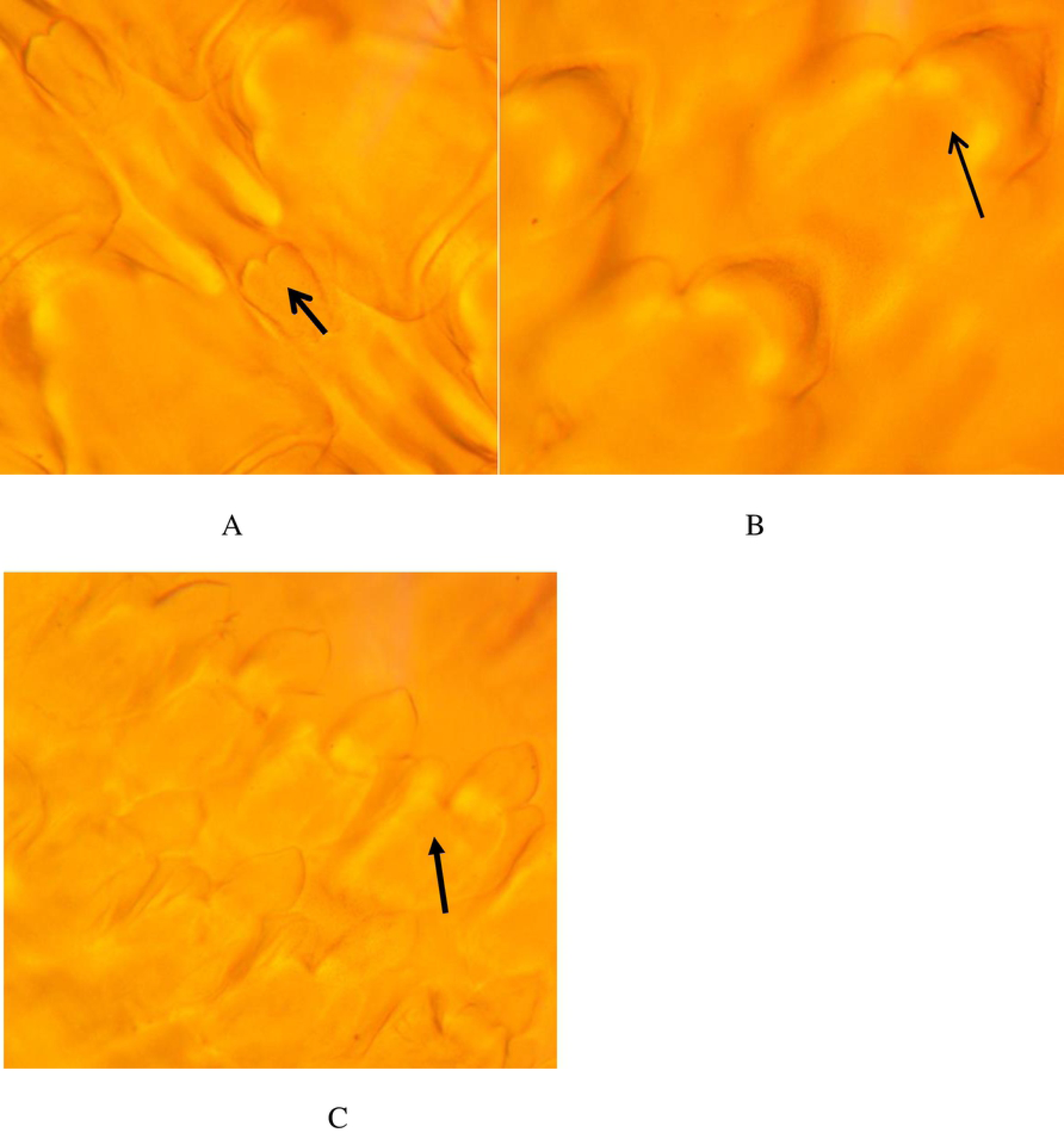
Apertural and abaperural views of shell of *Achatina fulica* collected From Itori, Ogun State, Nigeria. Scale bar = 1.0cm.

**Fig 3.**
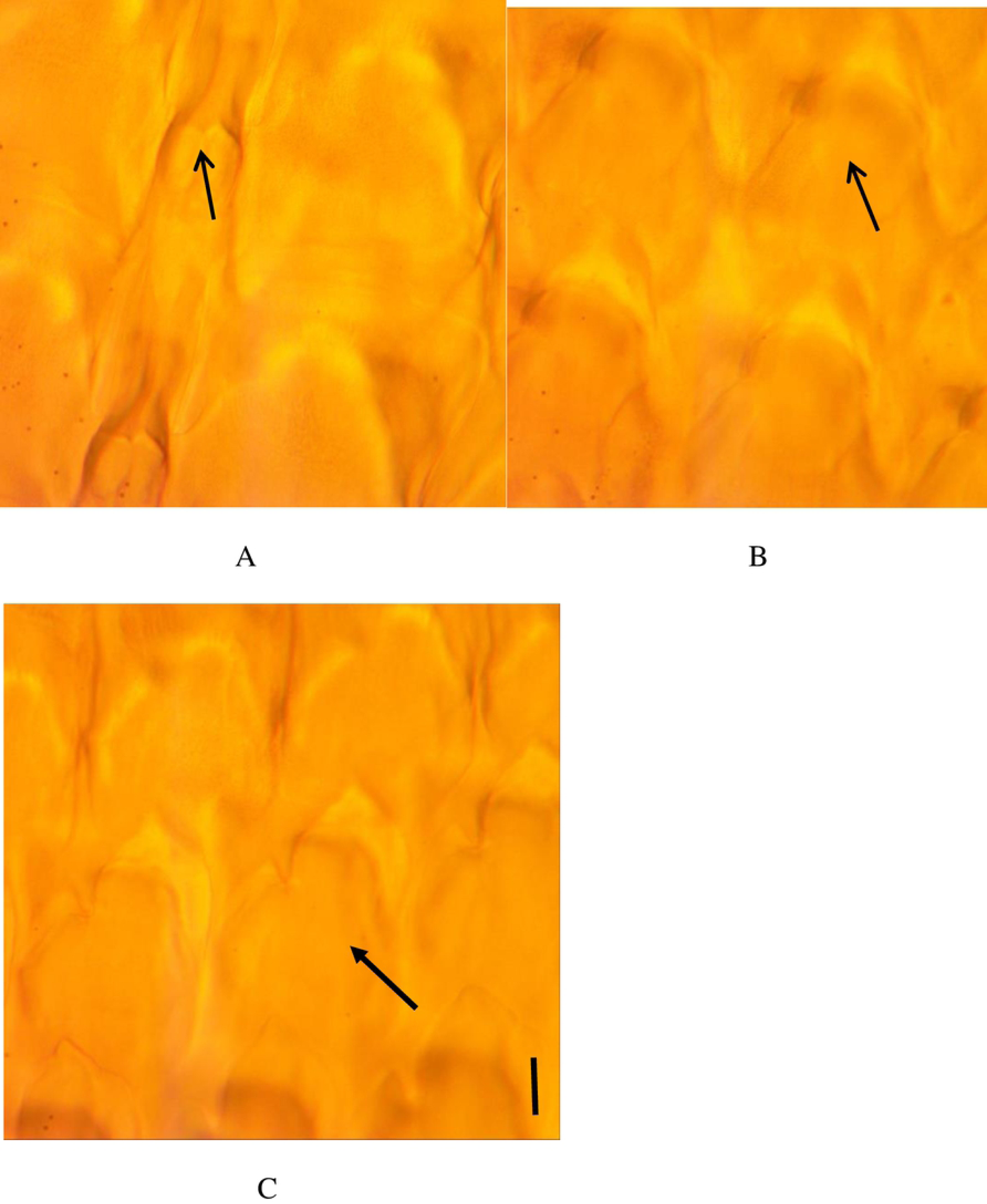
Apertural and abaperural views of shell of *Achatina reticulata* collected from Ibadan. Scale bar = 1.0 cm.

**Fig 4.**
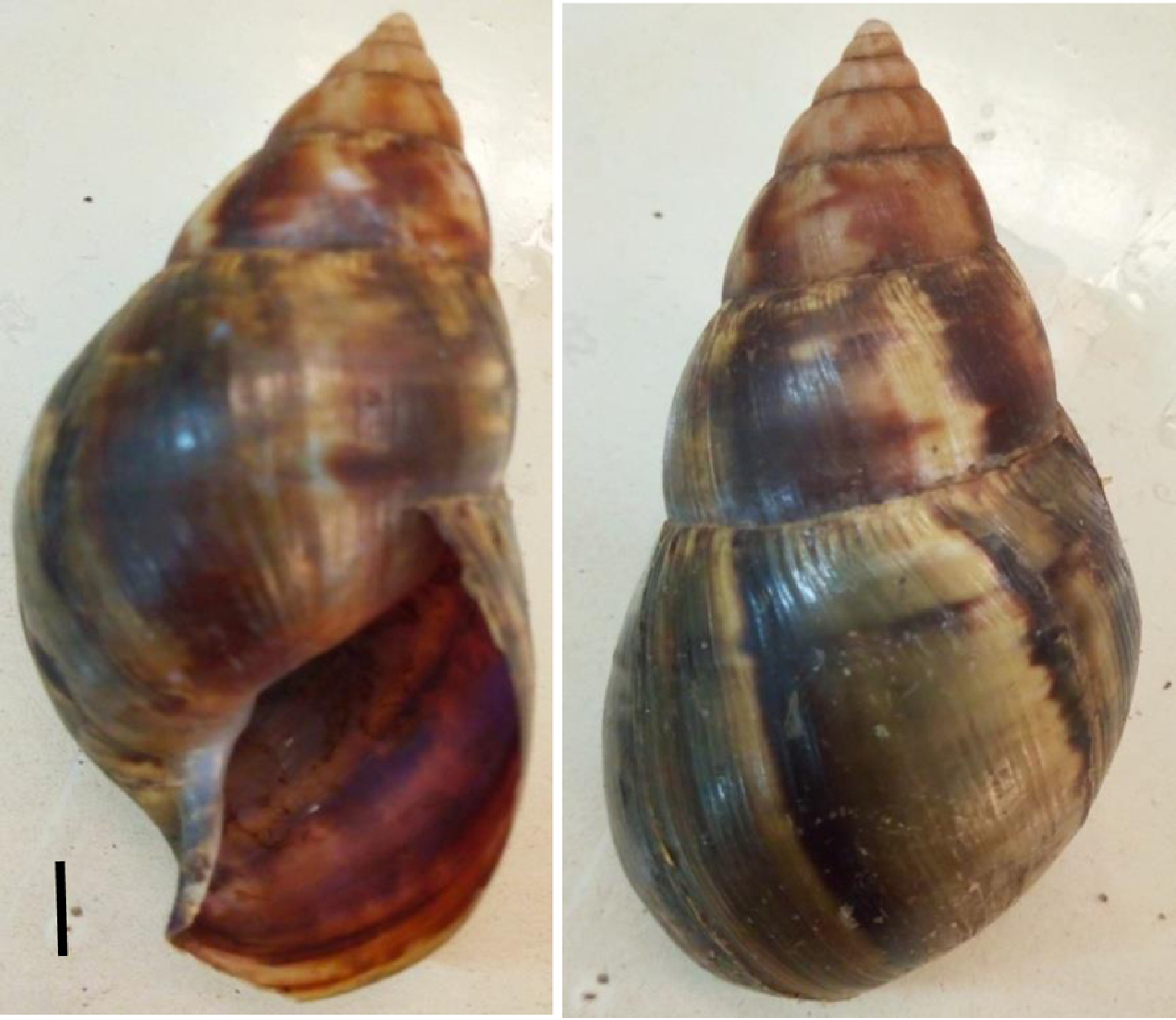
Apertural and abaperural views of shell of *Achatina reticulata* collected from Ibadan. Scale bar = 1.0 cm.

The three invasive species exhibited considerable morphometric similarities “Table 2”. They were high-spired species (shell height> shell width), and the shell spired index (SH/SW) ranged from1.8-2.0. The shell of *A. fulica* was most slender (SH/SW=2.0±0.04) followed by the shell of the shell of *A. reticulata* (SH/SW=2.0±0.1) and *A. albopicta* (SH/SW=1.8±0.1). However, the mean values of the shell parameters were significantly (p<0.05) larger for *A. reticulata* than *A. fulica* and *A. albopicta* “Table 2”. The shells of *Achatina reticulata* had a mean height of 134.0.3± 17.4 mm and was larger than the shell of *A. fulica* (90.0 ± 2.7 mm) and A. *albopicta* (104.1± 7.6 mm) “Table 2”. The aperture index (AP/AW) ranged from 1.9-2.3 among the three species. The shell shapes fit into the patterns described for invasive achatinidae [16]; this study showed that the three species had elongated spire with narrow body whorl and narrow aperture. *Achatina reticulata* was the largest of the three species in this study with conspicuous longitudinal sculptures on the body whorl. The high shell-spired indices of the three species may account for the ease with which they burrow or burry in the soil.

**Table 2.** Morphometric characterization of invasive Achatinidae collected from southwestern Nigeria.

| Character/Index | *Achatina fulica* | *Achatina albopicta* | *Achatina reticulata* |  |
| --- | --- | --- | --- | --- |
|  | X±SD(Range)<br>mm | X±SD (Range)<br>mm | X±SD (Range)<br>mm | P value |
| No. of whorls | 8.3±0.1(8.25-8.5) | 8.3±0.1(8.25-8.5) | 8.4±1.4 (8.25-8.5) | F=0.303;p=0.741<sup>a</sup> |
| Shell height(SH) | 90.0±2.7(87-93) | 104.1±7.6(98-120) | 134.0±17.4 (103-145) | F= 200.1;p=0.0001 |
| Shell width(SW) | 44.4±0.8(43-45) | 56.4±4.4(52-65) | 70.6±3.0 (69-75) | F=374.4;p=0.0000 |
| Spire height(SpH) | 27.4±2.5(25-31) | 30.5±1.3(29-32) | 41.0±1.0(40-42) | F=54.0;p=0.0000 |
| Body whorl height(BwH) | 57.3±3.9(52-64) | 77.0±5.8(70-85) | 88.8±2.3(88-92) | F=0.0949;p=0.9098<sup>a</sup> |
| Aperture height(AH) | 44.4±0.7(43-45) | 53.1±3.2(50-58) | 56.0±1.2(55-58) | F=104.8;p=0.0000 |
| Aperture width(AW) | 22.0±1.2(20-24) | 23.0±2.4(21-28) | 30.6±1.5(29-33) | F=64.0;p=0.0000 |
| SH/SW | 2.0±0.1(2.0-2.1) | 1.8±0.1(1.8-1.9) | 2.0±0.1(1.9-2.0) | F=27.1;p=0.0000 |
| AH/AW | 2.0±0.1(1.9-2.1) | 2.3±0.1(2.1-2.5) | 1.9±0.1(1.8-2.0) | F=28.9;p=0.0000 |
| BwH/SH | 1.3±0.1(1.2-1.5) | 2.3±0.1(2.1-2.5) | 1.9±0.1(1.8-2.0) | F=1.157;p=0.335<sup>a</sup> |
| SpH/BwH | 0.5±0.0(0.4-0.5) | 0.4±0.0(0.4-0.4) | 0.5±0.0(0.5-0.5) | F=41.1;p=0.0000 |

<sup>a</sup>Non-significant difference between mean values

The genitalia of three species were very identical, the basal genital structures were similar; the penes were slender and completely enclosed by the penial sheath “Fig 5”. The length of the penes varied from 10 mm to 12 mm in the three species “Table 3”. The vas deferens, free oviduct and the spermatheca duct were very long. The basal uterus is slightly greenish and the apical uterus is yellowish to pale cream. The hermaphroditic duct is highly convoluted and the basal part is pale cream in colour while the apical part is black in colour. The major differences in the basal genital structures are shown in “Table 3”. The genital structures were significantly different from the genital structures of the West African Achatinidae [17].

**Table 3.** Main features of the basal genital structures of *Achatina fulica, Achatina albopicta* and *Achatina reticulata* collected from southwestern Nigeria.

| Basal Genital structures | *Achatina fulica* | *Achatina albopicta* | *Achatina reticulata* |
| --- | --- | --- | --- |
| Penis | 10-11mm long | 10-11 mm long | 11-12 mm long |
| Penis sheath | Completely encloses the penis; basal portion of penis is swollen | Completely encloses the penis, swollen at the extreme end joining the basal vagina | Completely encloses the penis |
| Basal vas deferens | Emerges from the sheath almost at the upper end of sheath | Emerges from the sheath at the upper ¼ of the sheath | Emerges from the sheath close to the middle of the sheath |
| Basal vagina | Swollen and corm-like | Swollen and corm-like | Swollen and corm-like |

**Fig 5.**
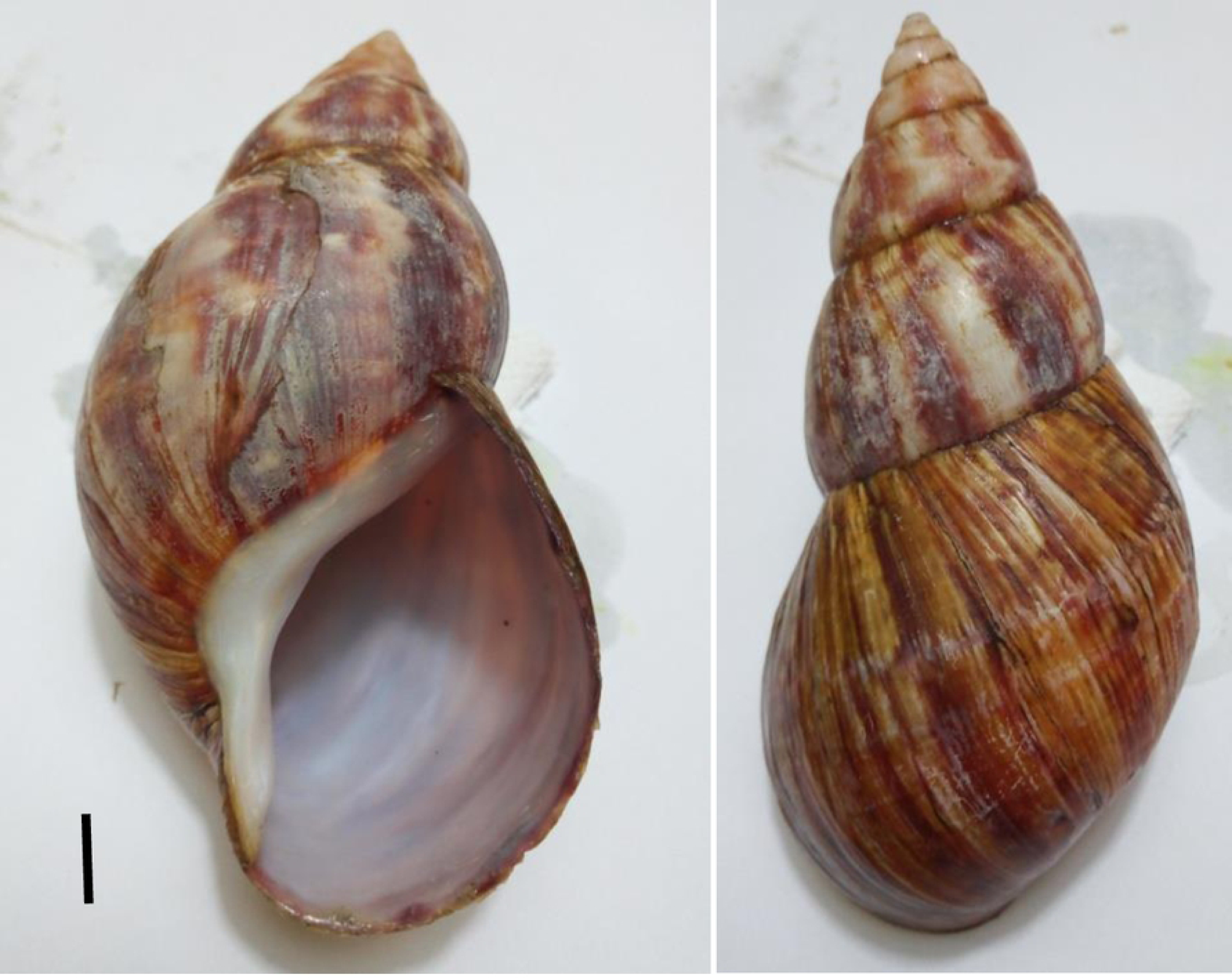
Diagrammatic illustrations of the basal portion of the genital system of *Achatina albopicta* (A), *Achatina reticulata* (B), *Achatina fulica* (C), showing the basal portion of uterus and prostrate (a), spermatheca (b), spermatheca duct, basal portion of the vagina (d), penis (e), basal portion of vas deferens (f) and the flagellum (g). Scale bar = 1.0 cm

The radulae of the three species followed the basic pattern for the pulmonata. There were a set of teeth on either side of the median tooth, the lateral teeth and another set of marginal teeth. The median teeth in the three species were very small compared to the lateral and marginal teeth and they were bicuspid. The shape of the median tooth differed in the three species “Table 4”. The lateral teeth of the three species were usually tricuspid, with a large centrally located cusp (mesocone) and two poorly developed accessory cusps(endocone and ectocone) on both sides of the central cusp. However, the first lateral tooth on the right side of the median tooth of *Achatina reticulata* “Fig 6” had well developed accessory cusps (endocone) next to the median tooth; the ectocone was not conspicuous. The radulae could be differentiated by the structures of their median teeth “Figs 6A, 7A, and 8A”. The accessory cusps, increased in number and levels of development from the first lateral tooth toward the margin, in the three species. The marginal teeth of the three species “Figs 6C, 7C and 8C” were characterized by larger number of cusps, with the mesocone and the accessory cusps (ectocone and endocone) further divided into smaller cusps. This study appears to be the first to present a detail study of the radula structures of the three invasive species. The radulae differed in the shape of median teeth, the development of the accessory cusps, particularly the endocone on the first lateral teeth.

**Table 4.** Main features of the radulae of *Achatina fulica, Achatina albopicta* and *Achatina reticulata* collected from southwestern Nigeria.

| Radula structures | *Achatina fulica* | *Achatina albopicta* | *Achatina reticulata* |
| --- | --- | --- | --- |
| Median tooth | Bicuspid with two unequal cusps, the larger cusp was pointed and taller than the smaller one with flat surface | Bicuspid and the two cusps are nearly of the same heights and sizes, with flat surfaces. | Bicuspid with two unequal cusps, the smaller cusp was pointed and taller than the larger cusp. The median tooth was almost obscured by the well-developed ectocone of the first right lateral tooth. |
| Lateral teeth | Tricuspid with well-developed mesocone | Tricuspid with well-developed serrated mesocone | Tricuspid with well-developed mesocone. The ectocone of the first right lateral tooth was well developed. |
| Marginal teeth | Numerous with the ectocone and endocone further divided into smaller cusps | Numerous with the ectocone further divided into smaller cusps | Numerous with the ectocone further divided into smaller cusps and the endocone greatly reduced. |

**Fig 6.**
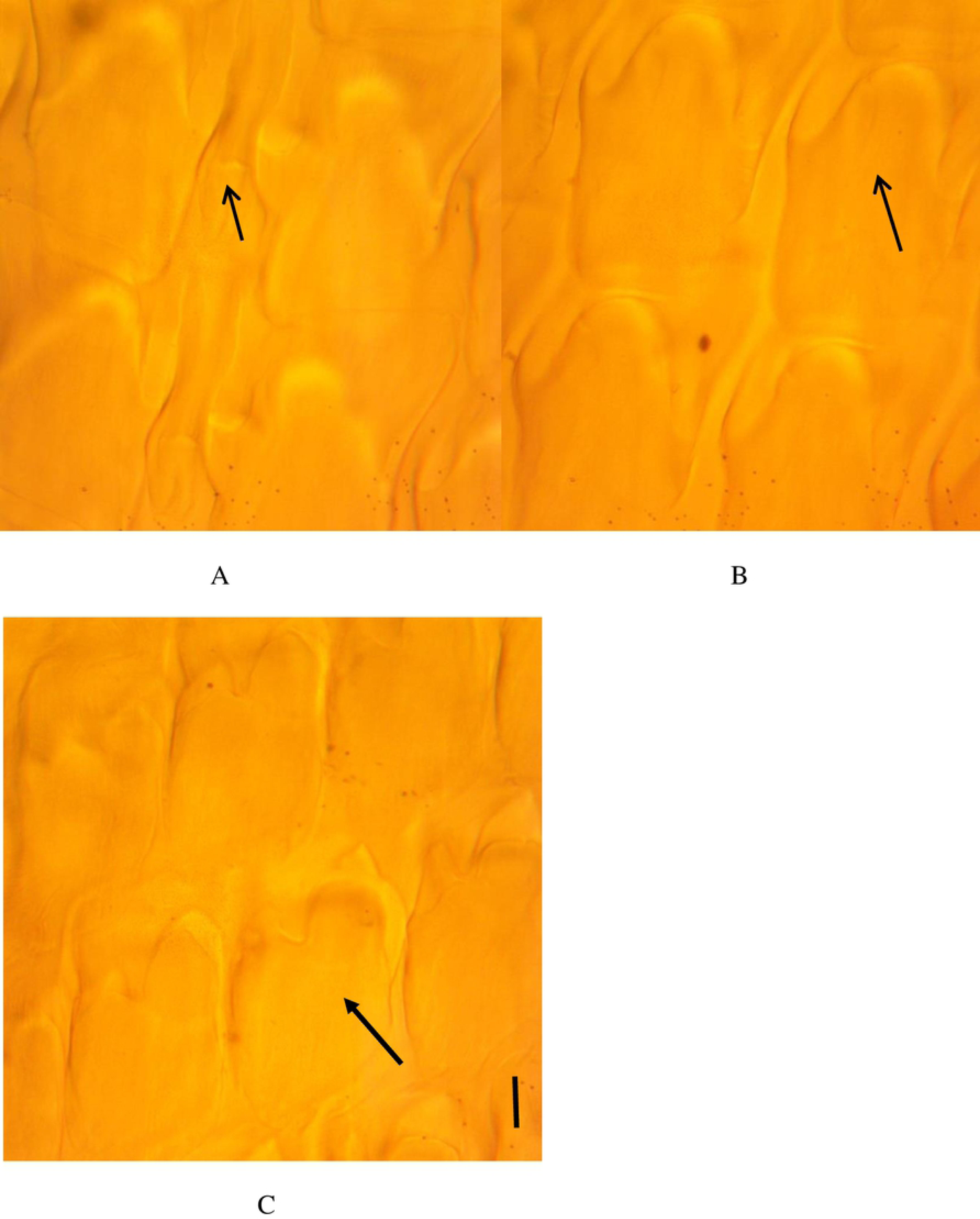
Micrograph of *Achatina reticulata* radula showing the bicuspid median tooth and the first right lateral teeth with well-developed endocone (A), tricuspid lateral teeth (B) and the marginal teeth (C). Scale bar = 1 μm.

**Fig 7.**
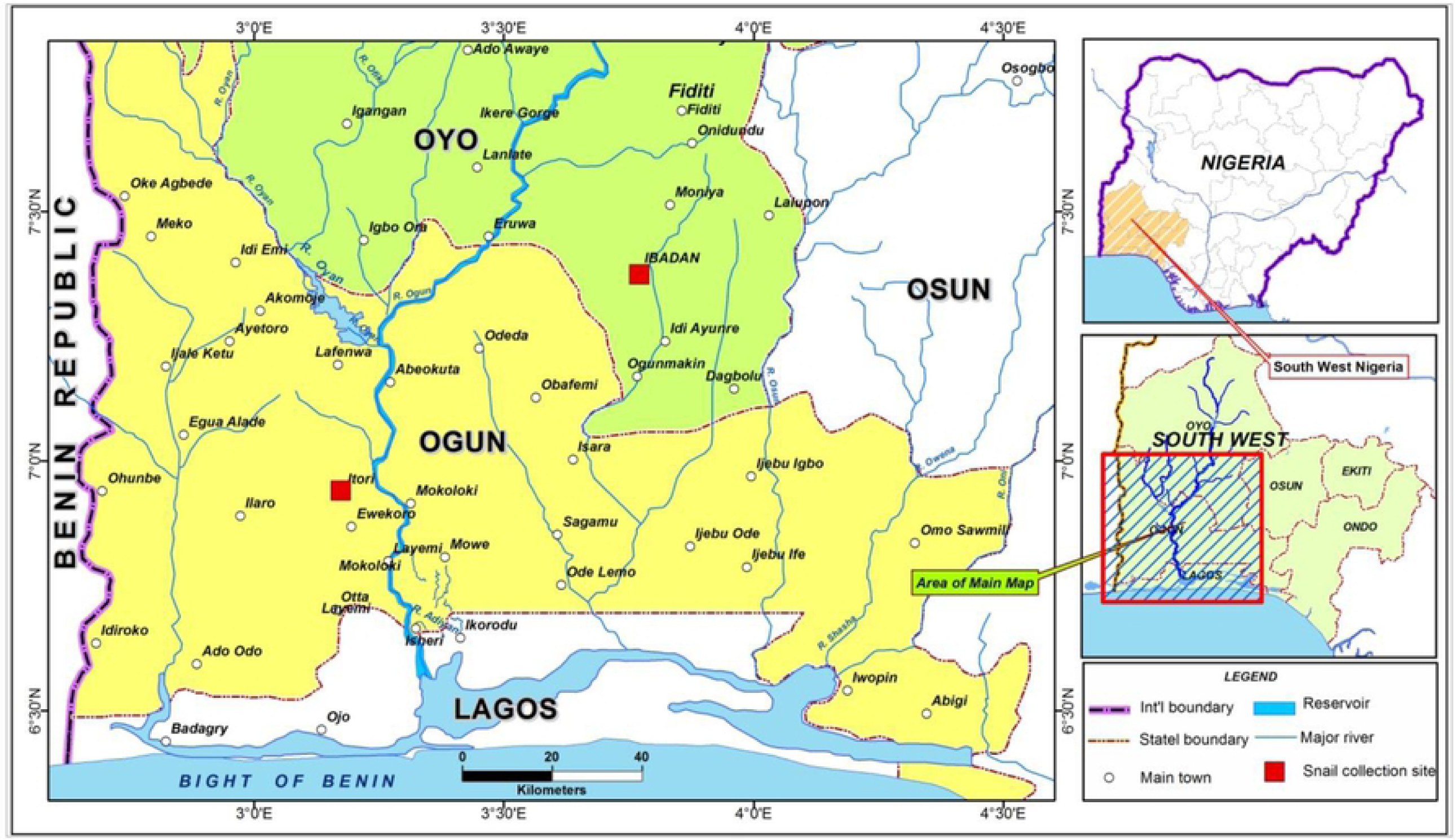
Micrograph of *Achatina albopicta* radula showing bicuspid median teeth (A), tricuspid lateral teeth and a serrated mesocone and the marginal teeth (C). Scale bar = 1 μm

**Fig 8.**
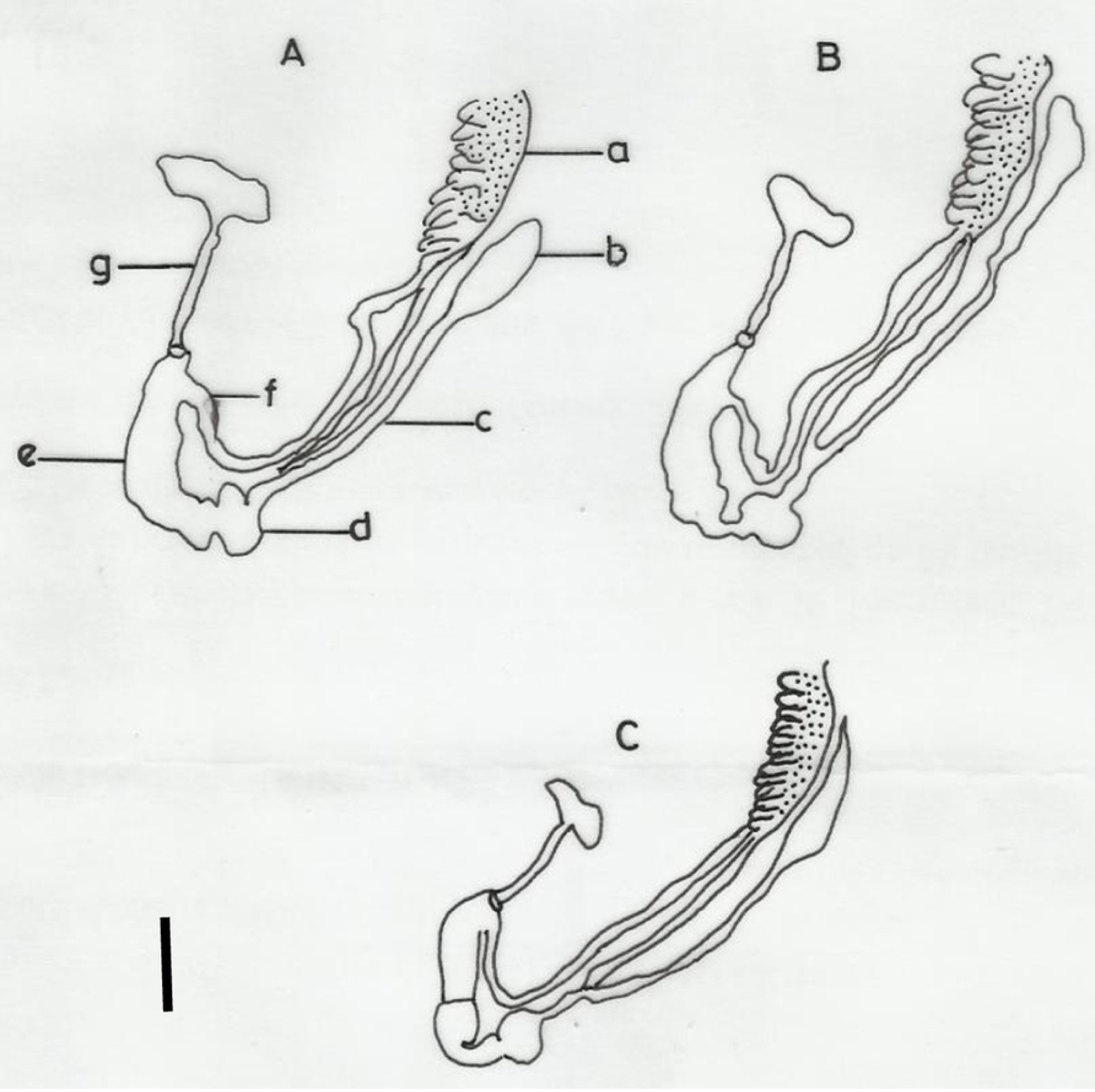
Micrograph of *Achatina fulica* radula showing bicuspid median teeth (A), tricuspid lateral teeth (B) and the marginal teeth (C). Scale bar = 1μm

## Conclusions

This study has shown that the invasive achatinids have become established in Nigeria and apart from *A. fulica*, other new subspecies *A. albopicta* and *A. reticulata* have been introduced. The reproductive structures, shell morphology and radula structures can be useful in the taxonomy of land snails. The study also suggests that the invasive species generally referred to as *A. fulica* in other areas, where they have been introduced, may be a mixture of subspecies of *Achatina* (subgenus *Lissachatina*) which were probably introduced together. It is therefore, likely that the other subspecies of the subgenus *Lissachatina* may be more widely spread outside East Africa than is currently documented due to misidentification.

## Acknowledgments

We thank Chris Odeh and TundeDisu, both of the Department of Zoology, University of Ibadan, Ibadan, Nigeria for their technical assistance. We are also grateful to Mr. Femi Balogun of the Department of Archaeology and Anthropology, University of Ibadan, Ibadan, Nigeria for

## Supporting information

S1 Table. Shell morphometrics of three invasive Achatinidae in southwest Nigeria

